# The periphery and the core properties explain the omnigenic model in the human interactome

**DOI:** 10.1101/749358

**Authors:** Bingbo Wang, Kimberly Glass, Annika Röhl, Marc Santolini, Damien C. Croteau-Chonka, Scott T. Weiss, Benjamin A. Raby, Amitabh Sharma

**Affiliations:** School of Computer Science and Technology, Xidian University, Xi’an 710071, China.; Center for Complex Network Research, Northeastern University, Boston, MA, USA; Channing Division of Network Medicine, Department of Medicine, Brigham and Women’s Hospital and Harvard Medical School, Boston, MA, USA; The Center for Research and Interdisciplinarity, Paris, France 75004

## Abstract

Understanding the connectivity patterns of genes in a localized disease neighborhood or disease module in a molecular interaction network (interactome) is a key step toward advancing the knowledge about molecular mechanisms underlying a complex disease. In this work, we introduce a framework that detects *peripheral* and *core* regions of a disease in the human interactome. We leverage gene expression data on 104 diseases and analyze the connectivity of differentially expressed genes (quantified by a p-value < 0.05) and their topological membership in the network to distinguish between peripheral and core genes. Per definition, peripheral and core genes are topologically different and we show that they also differ biologically. Core genes are more enriched for Genome-wide association study (GWAS) and Online Mendelian Inheritance in Man (OMIM) data, whereas peripheral genes are more shared across different disease states and their overlap helps predict disease proximity in the human interactome. Based on this observation, we propose a *flower* model to explain the organization of genes in the human interactome, with core genes of different diseases as the petals and the peripheral genes as the (shared) stem. We show that this network model is an important step towards finding novel drug targets and improving disease classification. Overall, we were able to demonstrate how perturbations percolate through the human interactome and contribute to peripheral and core regions, an important novel feature of the omnigenic model.

## Introduction

For complex disease phenotypes, genetic signals are spread across the genome and include genes without an obvious connection to pathobiology^1^. However, multiple studies have proposed that the genes associated with a disease are localized within a specific neighborhood of the human interactome (molecular interaction network)^2,3^, suggesting the existence of a disease neighborhood, or a connected subnetwork mechanistically linked to a disease phenotype ^3,4^. Moreover, genes, especially differentially expressed genes, exhibit a tendency for their protein products to interact with one another and agglomerate to form a connected component in the interactome^2,4–8^. If this is the case, then any gene whose expression is modified in a disease-relevant tissue is likely to be just a few steps away from one or more of the genes that have the strongest effects in a disease, defined as *core genes*^1^. It has been proposed that all genes expressed in disease relevant cells influence the function of the core genes^1^. Thus, the genes that are *a few steps* away from the core might be defined as the *peripheral part* of the disease neighborhood. Hence the critical genetic features of complex diseases might lie outside the core gene pathways in the peripheral portion of the gene network that might play a significant role in disease expression^9^. This prompts us to ask how the interactome can be explored to identify peripheral and core regions for complex traits and whether it is possible to identify the contribution of peripheral genes to infer disease-disease interactions.

We hypothesized that the conceptual distinction between core genes and peripheral genes in the human interactome will be useful for understanding human disease phenotypes. In this study, we attempt to explore the *component property* (Fig. 1) of the disease neighborhood that might help in explaining the core and peripheral properties of diseases in the human interactome. We develop a network-based framework that distinguishes the core and peripheral components of a given disease based on the local maxima of connectivity significance between the differentially expressed genes. The proposed network model helps to distinguish core and peripheral components by utilizing 184 gene expression data sets for 104 distinct disease phenotypes. Next, we characterize the core and peripheral genes, and explore their relative contribution to disease etiology. We provide evidence that genes comprising the core of a disease are disease-specific, while those in the periphery are shared among related disease conditions. This suggests a *flower-like structure* where core genes are petals and peripheral genes are the stem (Fig. 1). This flower structure helps in finding the commonalities between diseases and also explaining disease-disease relationships, which eventually results in discovering common biomarkers and therapeutic targets.

**Figure 1.**
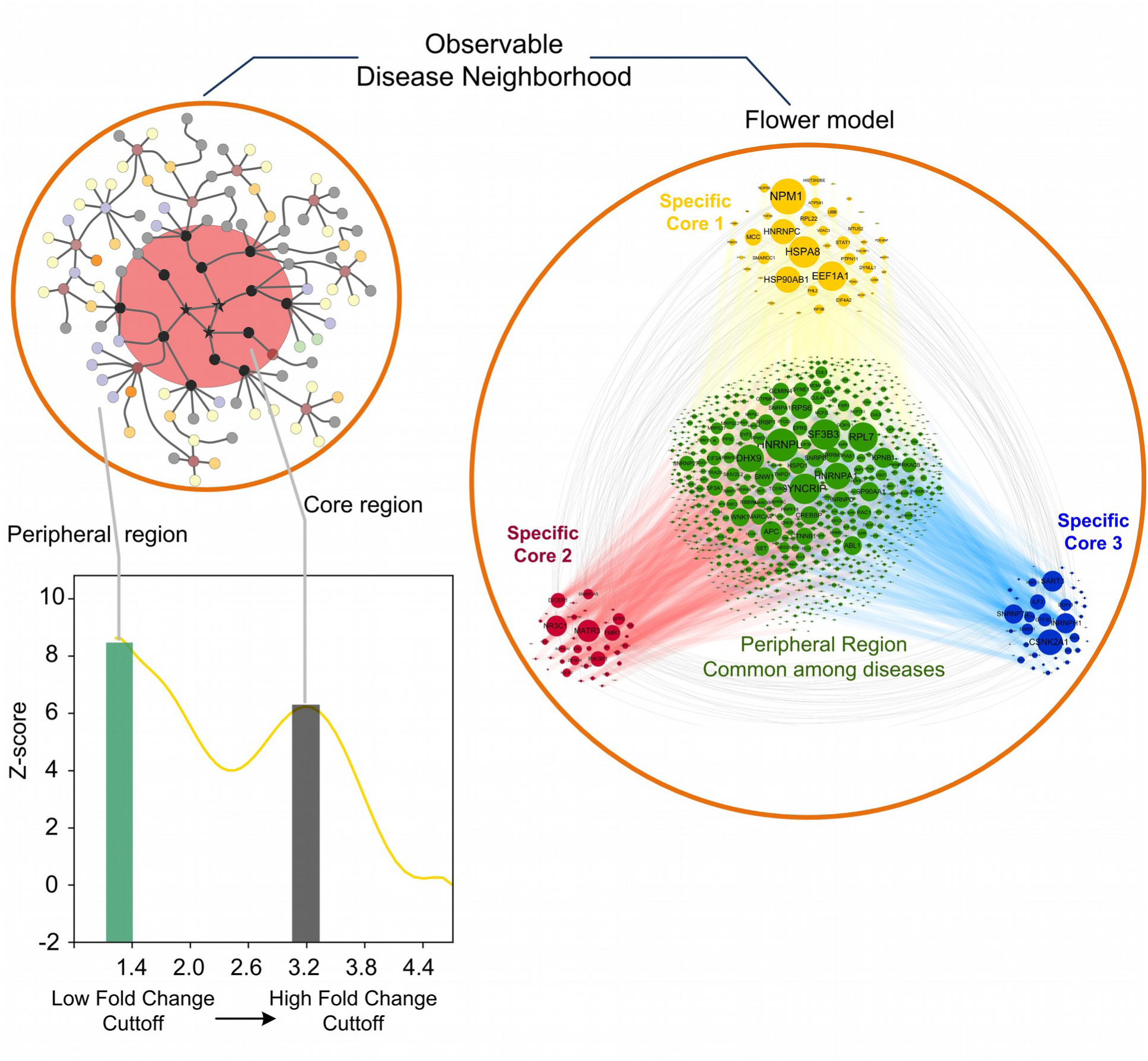
Overview of the approach to detect peripheral and core regions in the human interactome. A disease neighborhood consists of peripheral and core regions. While cores typically consist of genes specific to the underlying disease, peripheral genes are pleotropic and overlap among different diseases. This relational property can be described in a flower model. The yellow curve shows the differentially expressed genes fold change values vs. the z-scores. We can identify two peaks in this curve, where the LCC related to the first peak describes the peripheral disease neighborhood, while the core of this neighborhood can be found at the second peak.

## Results

### Identification of peripheral and core regions

We downloaded 184 gene expression data sets from Gene Expression Omnibus (GEO)^10^ that had clearly been defined as cases and controls for each disease (Supplementary note 1). Next, we applied limma^11^ to quantify the differential expression between cases and controls, resulting in a p-value and fold-change in the 184 data sets. Further, we selected the differentially expressed genes (DEGs) with p-value < 0.05 (unadjusted) for each data set and computed the largest connected component (LCC) induced by the DEGs for each disease. At different fold-change cutoffs, we selected the subsets of DEGs and identified the induced LCC that determines the disease neighborhood for the subset of DEGs (Fig. 1 and 2). To test, whether the LCC represents a non-random aggregation of genes, we compared the size of the LCC with the same number of random genes placed on the interactome and computed the LCC’s z-score (Supplementary note 3). Further, we applied three criteria to identify the core and peripheral regions: (1) the z-score of the LCC induced by the core genes need to be greater than 1.6, (2) the z-score of the LCC induced by the peripheral genes need to be greater than 1.6, and (3) the detectable ratio *d_ratio_* needs to be greater than 0.1. The *d_ratio_* quantifies the amplitude ratio of the second local maximum peak at the fold-change cutoff in the least squares polynomial fitting curve. The average *d_ratio_* for randomized data sets obtained by shuffling the cases and controls in gene expression data was 0.09 and we opted for *d_ratio_*>0.1 as an above average value. With the above criteria, we observed two peaks in the curve of the z-score values representing *local maxima* of DEGs aggregation in 135 from the total 184 data sets and the corresponding fold-change cutoffs were defined as *fc^low^* and *fc^high^* (Fig. 2a and Fig. S1). For example, for the pulmonary hypertension (GSE703) gene expression data, we identified two peaks at *fc^low^* =1.18 and *fc^high^* =2.09. At *fc^low^*, the significant LCC with z-score=8.81 consisted of 631 genes. While at threshold *fc^high^*, another significant LCC with z-score=6.67 had 62 genes. The two peaks suggest that for each disease, the disease neighborhood consisted of two distinct regions: a bigger peripheral region at *fc^low^* and a small core region at *fc^high^* (Fig. 1, 2a).

**Figure 2.**
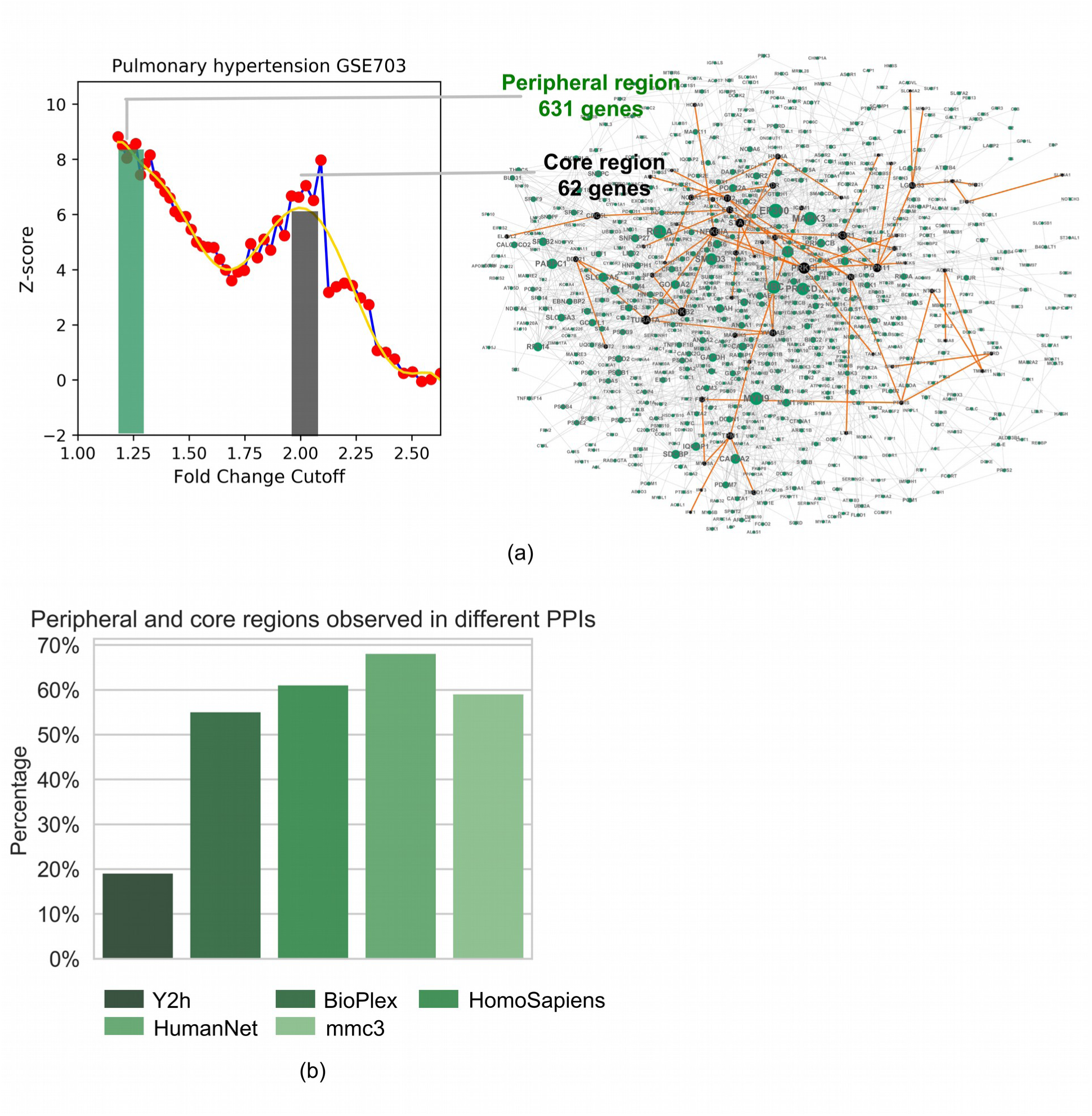
Application of the framework to 135 gene expression data sets. **(a)** Gene expression data of pulmonary hypertension illustrate how the framework identifies the core and peripheral regions. The red dots illustrate the LCC z-scores versus the fold change cutoffs and the yellow curve is the least square fit of these points. Two peaks can be recognized: one at *fc^low^* =1.18 and the other at *fc^high^* =2.09. The disease neighborhood is given by the LCC induced by the DEGs with a fold change larger than *fc^low^* and consists of 631 genes. The LCC induced by the DEGs with a fold change larger than *fc^high^* is the core, which consists of 62 highly fold change genes. **(b)** We tested the 135 data sets in five different protein-protein interaction networks: HuRI, Bioplex 2.0 (BioPlex), HINT (HomoSapiens), HumanNet.v1, and Mass spec (mmc3). The bar chart shows the percentage of the diseases for which observed the core and peripheral patterns.

We evaluated the robustness of local maxima pattern in different types of molecular interaction networks: (1) HuRI^12^ (http://interactome.baderlab.org/), (2) Bioplex 2.0^13^, (3) Homosapiens binary from HINT database^14^, (4) Masspec protein interaction data from Hein et. al 2015^15^, and (5) HumanNet functional network^16^ (see Methods). With the z-score greater than 1.6 for both LCCs at *fc^low^* and *fc^high^*, the peripheral and core neighborhood pattern was observed in more than 60% of the gene expression data sets in binary interactions from HINT, Massspec protein interactions, and HumanNet (Fig. 2b). With a higher z-score (>2), the pattern was still captured in more than 50% of the binary interaction data from HINT and HumanNet (SM Fig. S2a). Overall, these results indicate the peripheral and core patterns are a general phenomenon in both physical and functional human interaction networks.

Furthermore, when we shuffle the case and control labels in gene expression data and performed the differential expression analysis (Fig. S1), the *d_ratio_* values were reduced from an average of 0.32 to 0.09 and the core region at *fc^high^* vanished in the shuffled differential expression data. This indicate that the core patterns are specific to the disease perturbations in the gene expression data.

### Topological characteristics of core and peripheral genes

Topological analysis of the core and the periphery of the human interactome indicated that core genes have a significantly higher degree, betweenness, and closeness centralities ^17^ compared to peripheral genes, with the exception of the clustering coefficient^18^ (Fig. 3a). We used the Wilcoxon signed-rank^33^ test to calculate the p-values for these comparisons (see Methods). This suggests that the core genes with high fold-changes tend to be t*opologically central* in the disease neighborhood, while the peripheral genes have low degree compared to core with low fold-change. Furthermore, we found that removing the core has little impact in fragmenting the disease neighborhood or the sub-network, but if we delete the first neighbors of the core genes defined as the dominant range (DR), it leads to a higher fragmentation of the disease sub-network (Fig. 3 b, SM Fig. S4a,b). This implies that the core genes maintain the structure of the disease neighborhood by their immediate neighbours in the interactome.

**Figure 3.**
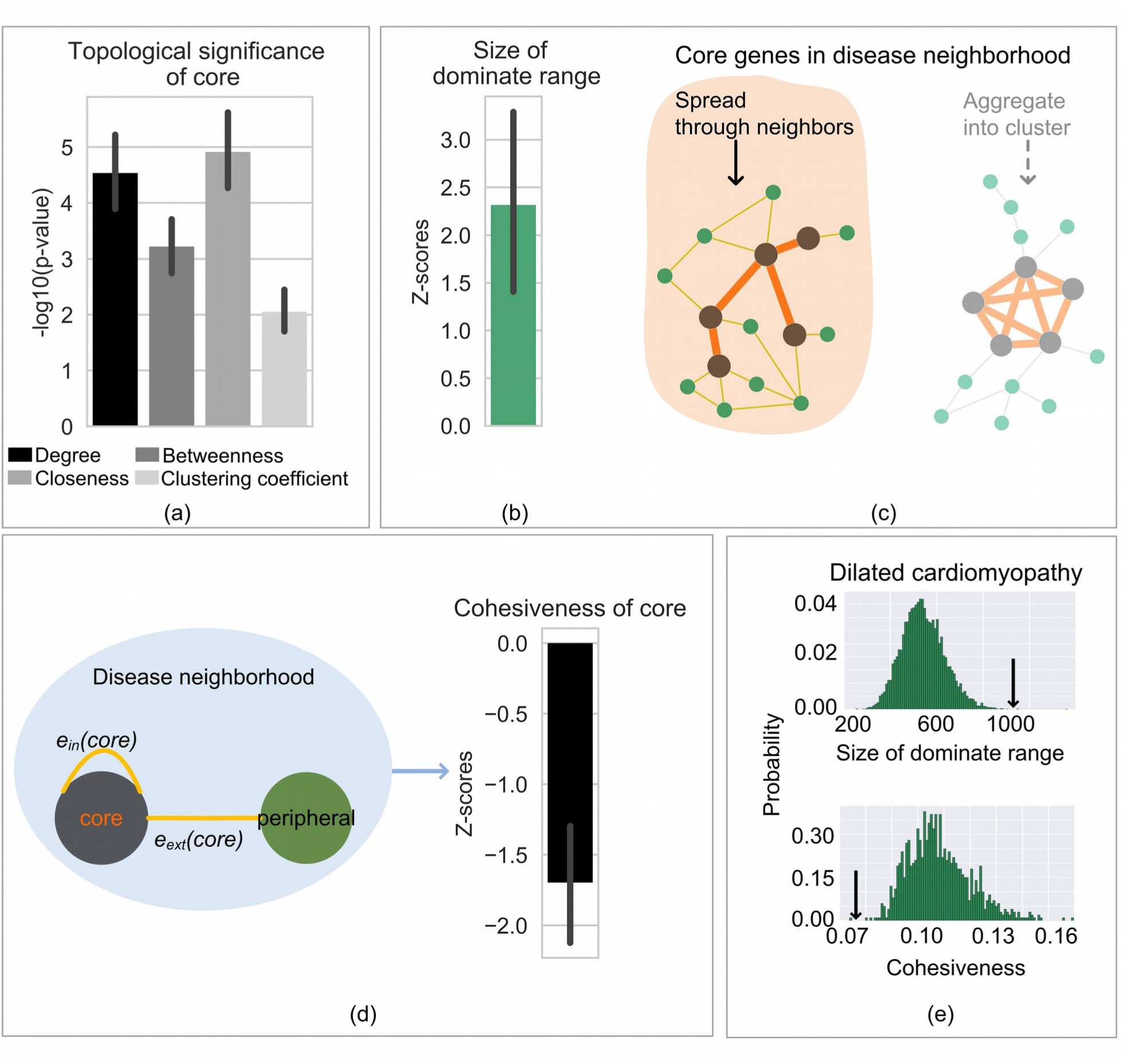
Comparison of the topological characteristics of the core and the peripheral regions. **(a)** Comparison of centrality measures between core and peripheral regions in 135 data sets. For each centrality index, a p-value of 0.05 or smaller indicates that the centrality values of the core genes are significantly different to the values of the peripheral genes. **(b)** The dominate range (DR) consists of the first neighbors of the core. A positive z-score indicates that a DR of a given core is larger than a DR given by a random set of nodes of the same size as the core (compared with 1,000 random counterparts) in each data set. The box plot shows all z-scores (most of them >=2) of these 137 studied data sets **(c)** The schema illustrates that the genes of a core rarely cluster, but rather tend to spread through neighbors into the disease neighborhood. **(d)** The figure illustrates cohesiveness of the core genes, where *e_in_(core),* resp. *e_ext_(core),* denotes the number of internal edges, resp. external edges of the core. For each data set, we compared the cohesiveness of the core with the 1,000 RCCs using genes from the periphery to get a z-score. The box plot shows the negative z-scores (<=-2 on average), indicating the significant small cohesiveness of the cores in the 135 studied disease neighborhoods. **(e)** Dilated cardiomyopathy expression data shows the significance of the size of the dominant range and its cohesiveness.

Further, we tested whether core genes aggregate into a dense community or if their connectivity is spread throughout the whole disease neighborhood of the human interactome (Fig. 3c) by using the notion of a cluster. A cluster represents a locally dense sub-network that has more internal than external connections^19^. In comparison, an outspread sub-network has less internal connections and is more spread out to connect with the adjacent neighbors. Thus, we quantified the extent to which the core corresponds to a topological cluster by the cohesiveness index, which is calculated by dividing the number of internal edges by the sum of the number of internal and external edges (see Methods). We calculated the cohesiveness of core genes and compared it with the cohesiveness index obtained for random connected components (RCCs) of the same size (Fig. 3d, 3e). Interestingly, we find that core genes have the same edge density as RCCs (SM Fig. S4c) and moreover, their cohesiveness values were significantly lower than RCCs (Fig. 3d), suggesting that the core genes do not tend to be part of a dense local cluster, but instead extend to communicate with other nodes in the disease neighborhood through the peripheral genes. Furthermore, the low clustering coefficient values confirm that core genes avoid dense clustering (Fig. 3a), an observation previously reported for the disease genes in the human interactome^20^.

### Biological significance of the core compared with the periphery in the disease neighborhood of the human interactome

To further differentiate the core and peripheral genes of diseases, we tested the enrichment of peripheral and core components in the intrinsically disease-causing disordered proteins (IDPs) ^21^ and disease genes annotated by GWAS^22^ and OMIM^23^. We combined the results from 135 data sets in the human interactome to create a unique global core (5,743 genes) and a unique global periphery (9,918 genes), where the genes of the global core and the global periphery are not overlapping (see Methods). The global core was highly enriched in IDPs (p-value=5.64e-24) and disease genes (p-value=4.95e-46) (Fig. 4a), while the global periphery showed no enrichment. For example, *IL2RB*^24^, *HLADRB1*^25^, *HLA-DPA1*^26^, *HLA-DPB1*^26^, *DCN*^27^, *CCL5*^28^, *TNF*^28^, *VEGFA*^29^, *CCR5*^30^, *CSF1R*^31^, *SLAMF1*^32^, and IL18RAP^33^ are genes that are known to be associated with asthma (GSE65204) that were part of the global core (SM Fig. S5a). The 84 asthma core genes were connected to 1,048 peripheral genes that might add a small effect to disease risk.

**Figure 4.**
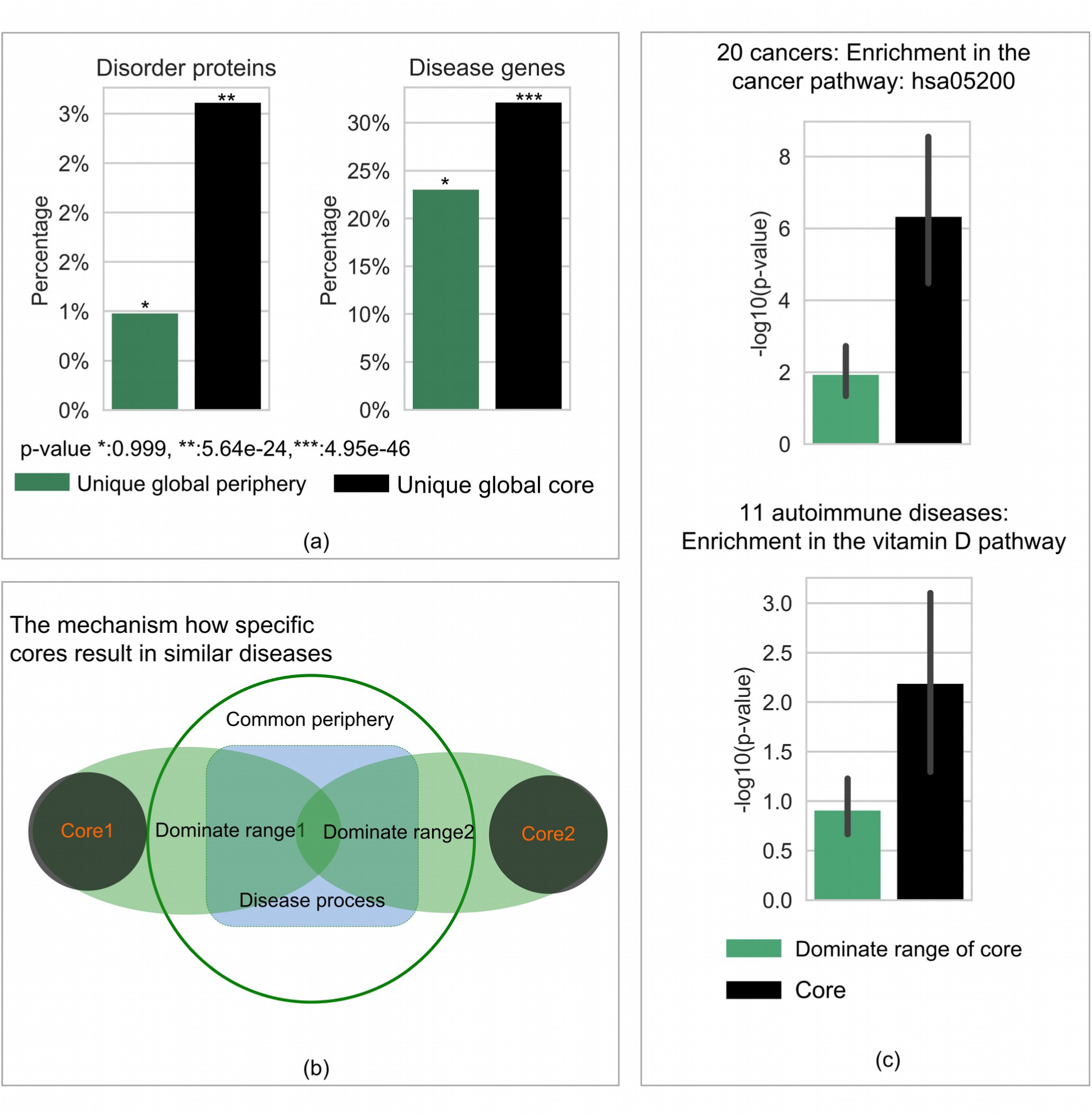
Biological significance of the periphery and the core regions. **(a)** The genes in the cores and peripheries from the 135 data sets were combined to form a unique global core and a unique global periphery. We observed more disease genes and intrinsically disordered proteins (IDPs) in the global core compared to the global periphery. **(b)** The mechanism through which cores influence biological pathways associated with a disease through the first neighbors (dominate range). **(c)** The box plot shows the significance of the enrichment of first neighbors for data sets associated with cancer and autoimmune diseases.

One key question that needs to be studied is the role of core genes in key disease pathways and biological processes of similar diseases in the human interactome. Since the immediate neighbors of the core might play key roles in the disease neighborhood, we assume that specific cores intervene in common processes of similar diseases through their immediate neighbors and are not clumped in disease-relevant pathways (Fig. 4b). We validated this hypothesis in 20 different types of cancer and 11 autoimmune diseases (Fig. 4c, see Methods and Supplementary Table 6): (1) For 20 different types of cancer, we considered the cancer pathways from the KEGG database (KEGG: hsa05200) as the common biological processes among these different types of cancer. We find a significant enrichment (Fig. 4c, p-value <=1.0e-4) of their first neighbors in the cancer processes and not the specific cores, thus suggesting that cores influence this common process through their immediate neighbors, ultimately leading to the disease phenotype. (2) Similarly, we observe the same phenomenon with 11 different autoimmune diseases. Since it is known that vitamin D plays an important role in auto-immune diseases^34^, we took the vitamin D pathway as their common process among these 11 diseases. The cores are rarely enriched for the vitamin D pathway, but through their immediate neighbors significantly affect this common process (Fig. 4c, p-values<=1.0e-2).

### Disease-disease relationship based on peripheral genes

Next, we tested genes overlap in the peripheral and core regions of the disease neighborhood (hypergeometric test). We observed a significant overlap between the peripheries of diseases (Fig. 5) and a non-significant overlap between the cores, indicating that the peripheral regions share more genes. Moreover, we observed a higher Jaccard similarity^35^ between pairs of peripheries compared to pairs of cores (Fig. 5b,d,e). This result holds true with other similarity measures (Sørensen–Dice coefficient, Simpson index, cosine index and geometric index^35^) (SM Fig. S6). Additionally, we found that the peripheral regions are significantly overlapping, where this does not hold true for the cores of different diseases (Fig. 5c). Further, we performed Hierarchical linkage-based clustering on the peripheries and cores based on their overlap and found that the cores (98.5%) are dissimilar while the peripheries (93.8%) have a high degree of similarity (Fig. 5e). For example, the relation between cardiomyopathy (GSE1869), Crohn’s disease (GSE3365), and dilated cardiomyopathy (GSE3586) disease neighborhood are shown in Fig. 5a. We assigned to each gene a three-dimensional coordinate (*fc_1_*, *fc_2_*, *fc_3_*) based on the fold-change level in the three data sets. The disjoint cores consisted of 40, 40, and 46 genes respectively, which were located at the end of each coordinate. There was a significant overlap (1,184 genes) between peripheries of similar diseases, e. g. cardiomyopathy and dilated cardiomyopathy are located at the centre. The three diseases had 767 overlapping genes in peripheries. These overlapping genes were enriched for inflammatory related pathways like TGF-beta signalling (Adj.p=0.0001), IL-6 signalling (Adj.p=0.002), and TNF-alpha NF-kB signalling (Adj.p=0.02) in ConsensusPathDB^36^. This is consistent with the crucial role inflammation plays in defining the two cardiac diseases^37^ and Crohn’s disease is a chronic inflammatory condition^38^. Thus, in the three-dimensional space, core and periphery form a flower-like structure with higher overlap of peripheral genes and a specific core for each disease as shown in Fig. 5a.

**Figure 5.**
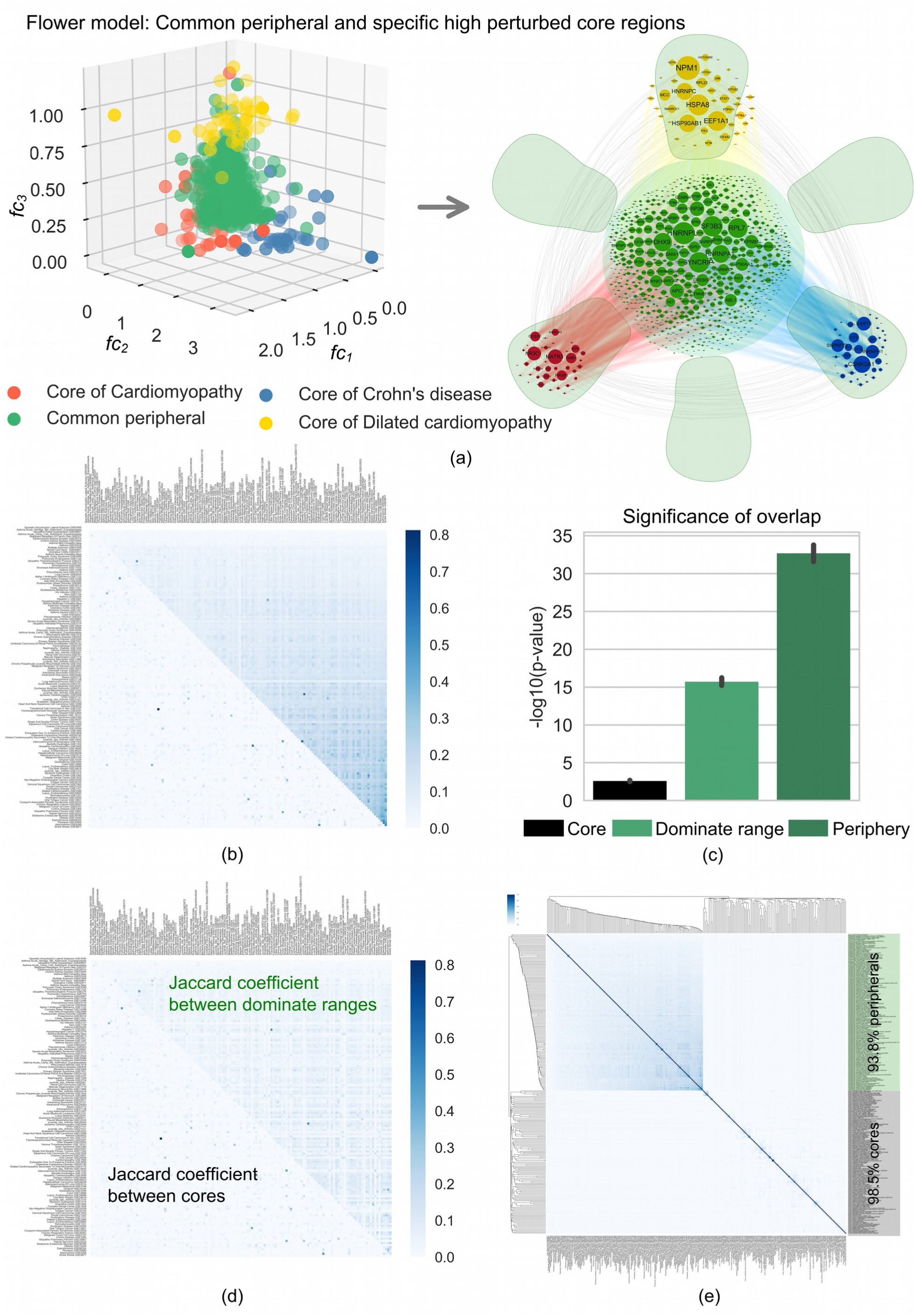
Flower model depicting peripheral as center and core regions as petals. **(a)** Left: Each axis represents the fold-change values of cardiomyopathy, Crohn’s disease, and dilated cardiomyopathy peripheral and core regions. The core genes with higher fold-change values are more specific to diseases and therefore are less overlapping compared to peripheral genes (green). Right: The flower like structure for the three diseases. The stem, highlighted in green, consists of the common genes from the peripheries. The color-coded petals present the specific cores. **(b)** The heat map represents the Jaccard coefficient values of the cores and the peripheral genes in 135 data sets. The lower triangle shows the smaller overlap between the different cores and the upper triangle shows the visible larger overlap between the peripheries. **(c)** The box plot shows the p-values for the overlap of the genes from the cores, dominate ranges (first neighbors of the core genes), and peripheries. **(d)** The heat map represents the Jaccard coefficients between cores and dominate ranges for 135 data sets. **(e)** Hierarchical linkage clustering on the peripheries and cores based on the overlap. We curated 135 cores and 135 peripheries together to form 270 gene sets, constructed a similarity matrix (270*270, based on the Jaccard coefficient) of these sets, and did hierarchical linkage clustering on this similarity matrix. 98.5% of the core sets are too specific to be clustered, and the higher similarity values concentrate on the 93.8% periphery sets.

To further test the utility of core and peripheral genes in the context of disease classification, we quantify the disease-disease relationships among 104 diseases by the connection specificity index (CSI)^35^ (see Methods) as shown in Fig. 6a. All 104 distinct diseases were grouped into 16 classes manually and are highlighted by different colors according to the medical subject heading (MeSH) terms in Fig. 6a (see Methods). Similarity values (link thickness is proportional to the CSI values) between intra-group diseases are significantly higher than values between inter-group diseases (p-value=7.72e-22), which indicates that similar diseases share the periphery and are cohesively connected to each other. In addition, this disease-disease network provides novel insight about disease-disease relationships. For example, pulmonary emphysema (GSE1122), alpha-1-antitrypsin deficiency (GSE1122), pulmonary hypertension (GSE703), and malignant neoplasm of the stomach (GSE2685) are tightly connected based on the periphery but are sparsely connected due to minimal core similarity (Fig. 6a). Alpha-1 antitrypsin deficiency is a hereditary disorder associated with pulmonary emphysema, inflammatory, autoimmune, and neoplastic diseases^39^. Further, we tested whether CSI relationships among diseases based on the peripheral proximity can also be used for drug repurposing among contiguous diseases in the human interactome (see Methods). We found that 61 diseases (out of 104 distinct diseases) had known approved drugs (The gold standard database, repoDB^40^) and/or drug targets (DrugBank^41^ database). We observed that 114 disease pairs had common drug targets in their common peripheral neighborhood. The box plot shows that the common drug targets are significantly enriched in the overlapping part of the periphery for these 114 disease pairs (hypergeometric p-value, taking the whole network as background) as shown in Fig. 6b. For example, for ankylosing spondylitis (GSE11886), rheumatoid arthritis (GSE1919), and Crohn’s disease (GSE24287), we found one common drug target, protein FCGR2C, and their common approved drugs, Etanercept (DB00005) and Adalimumab (DB00051), located in their overlapped peripheries (p-value=3.4e-05). For ankylosing spondylitis and rheumatoid arthritis, four common drug targets proteins (FCGR2A, FCGR2C, LTA and CA2) were targeted by three common approved drugs Adalimumab (DB00051), Etanercept (DB00005), and Celecoxib (DB00482), located in their overlapping peripheries (p-value=2.5e-04). Furthermore, for Crohn’s disease and rheumatoid arthritis, two common drug target proteins (FCGR2C and FCGR3A) and three common drugs Adalimumab (DB00051), Etanercept (DB00005), and Rituximab (DB00073) were found in their overlapped peripheries (p-value=0.009). This illustrates that the peripheral overlap can reveal more about disease-disease relationships and can be exploited for drug repurposing among similar diseases.

**Figure 6.**
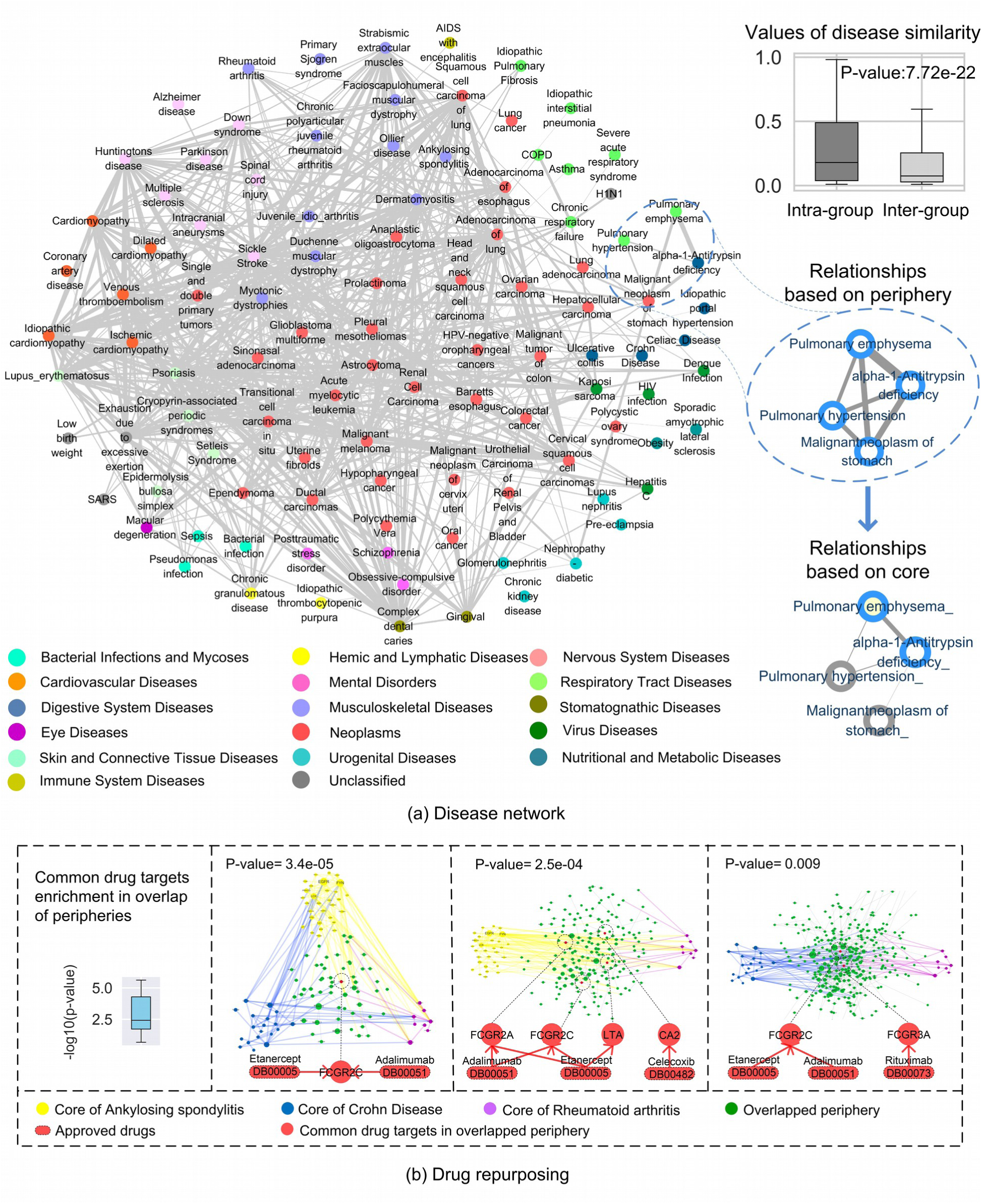
Disease-disease relationship based on the peripheral regions. **(a)** The disease-disease network of 104 distinct diseases constructed based on overlapping peripheries, where the 16 different disease groups (divided according to MeSH terms) are highlighted by different colors. Each node corresponds to a distinct disease, where the four gray nodes represent diseases that are unclassified in MeSH. Two diseases are connected by a link if they are related to each other and the thickness of the edge is proportional to the connection specificity index of the overlap of their peripheries. The box plot shows that the similarity values between intra-group diseases are significantly higher than between inter-groups diseases (p-value=7.72e-22). We highlight in the dotted box the strong relationship among pulmonary emphysema, alpha-1-antitrypsin deficiency, pulmonary hypertension, and malignant neoplasm of stomach, while their relationships were weak based on the overlap of their cores. **(b)** The box plot shows that common drug targets are significantly enriched in the overlapping part of the peripheries for 114 disease pairs. For example, the cores of ankylosing spondylitis, rheumatoid arthritis, and Crohn’s disease are highlighted by yellow, purple, and blue, respectively. The genes of the overlapping peripheries are highlighted in green. We identified five common drug targets (protein FCGR2C, FCGR2A, LTA, CA2 and FCGR3A) and four common approved drugs (Adalimumab (DB00051), Etanercept (DB00005), Celecoxib (DB00482), and Rituximab (DB00073)) among the overlapping genes in the peripheries. We highlighted the nodes and the directed links, representing approved drug-target relationships, in red.

## Discussion

An important feature of the omnigenic model, first articulated by Boyle et. al. (2017), was the classification of genes as *peripheral*, which are part of many diseases and thus are pleiotropic, and *core* genes, which are more specific for each disease^1^. In this work, we integrated differential gene expression with connectivity significance in the human interactome to capture peripheral and core disease regions by identifying the two *local maxima* of DEGs at different fold-changes. We characterized the peripheral and core gene regions of 104 distinct disease phenotypes. Furthermore, the *local maxima* of DEGs were robust in different biological networks. We observed that for these 104 diseases a core was comprised, on average, of 80 genes and surrounded by a peripheral region, consisting of 2,479 genes (Fig S3). The average fold change to discover the peripheral regions was 1.3 and it was 3.17 for discovering core regions in 135 gene expression data sets (Fig S3a). This illustrates that the small core of each disease contributes the strongest effects (high fold-change) to total disease association. However, these genes are also connected to numerous peripheral genes in the neighborhood, with mild to moderate modified expressions. Thus, it is not any individual gene that is likely to affect the function of the core genes as proposed by Boyle *et al.* (2017), but rather a set of genes that are significantly connected to the core and that show changes in their expression levels related to the disease. Furthermore, the topologically distinct nature of peripheral and core genes explain that perturbation in the high-degree core might lead to higher chances of disease susceptibility in the disease neighborhood, which is supported by the enrichment of core genes in the GWAS data. Moreover, the core mediates the propagation through the peripheral nodes in the disease neighborhood by avoiding the inner clustering.

The disease-disease network, which was constructed based on the similarity of peripheral regions, helps clustering and classifying the 104 distinct diseases as well as finding common therapeutic targets. The high similarity among the peripheral regions of diseases explains the key feature of the omnigenic model, which assumes that peripheral genes contribute to the risk of many diseases and are therefore likely to be pleiotropic. This observation is supported by the previous suggestions that pleiotropy is a common property of genes associated with disease traits ^42^. The flower-like structure of disease-disease relationships demonstrate that disease associated signals are not dispersed in the human interactome but rather that common pathways are altered and may spread to one or more diseases as shown in the example of cardiomyopathy, dilated cardiomyopathy, and Crohn’s disease. The core disease genes intervene in common processes of similar diseases through their immediate neighbors and are not directly enriched in disease-relevant pathways. We were able to establish this core property with 20 types of cancer and 11 autoimmune diseases. Moreover, it has been previously observed that highly diverse complex disease modules tend to overlap in the human interactome and are enriched with shared pathways like inflammation^43^.

Overall, we were able to demonstrate the recent hypothesis proposed by Boyle *et al.* (2017), that states that perturbations, which percolate through the molecular interaction system contributing to a disease risk, are driven by the large number of peripheral genes with no direct relevance to the disease and are propagated through a much smaller number of core genes, that have direct effects. The developed approach provides a framework that might help address numerous problems associated with disease gene identification, drug repurposing, and a general understanding of human complex diseases.

## Methods

### Gene expression data for model building

We collected 184 gene expression profiles from GEO for 104 diseases. Next, we applied the limma R package (Version 3.10.1)^11^ for differential expression analysis from the Bioconductor project^44,45^. We considered the genes with an unadjusted p-value < 0.05 as differentially expressed and selected them for further analysis (see SM Note 1.2 for more details).

### Mapping to the human Interactome

We constructed the PPI network using different sources: (1) Binary interactions from high-quality yeast-to-hybrid data sets; (2) Literature curated interactions typically obtained by low throughput experiments; (3) Kinase-substrate pairs, and (4) Signalling interactions. The union of all interactions yields a network of 16,461 proteins that are connected by 239,305 physical interactions (see SM Note 1.3 and Supplementary Table 4 for more details).

### Detection of the pattern

We start with differentially-expressed genes (DEGs) set*Ω*, and its subsets*Φ^k^*with different fold-change cutoffs *fc^k^* for each disease. Thus, we have

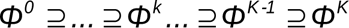

determined by

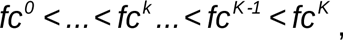

where *fc*^0^=*min*{*fc_i_* **∨** *g_i_ Ω*} with *fc_i_* being the fold-change value of gene *g_i_*. Furthermore, *fc^K^* =*max*{*fc_i_* **∨** *g_i_* ∈ *Ω*} and *Φ^K^* ={*g_i_* **∨** *fc_i_* ⩾ *fc^k^*, *g_i_* ∈ *Ω*}. Thus, *Φ^k^* contains all genes with a fold-change higher or equal to *fc^k^*.

To quantify the significance of the connectivity of *Φ*^0^, … *,Φ^k^*, …*, Φ^K^* we calculated their z-scores *z*^0^, …, *z^k^*, …, *z^K^*, respectively. To do so we computed LCC*^k^* which is the LCC induced by*_Φ_^k^*and next compared the size of LCC*^k^* with 1,000 LCCs given by sets of genes of the same size as*Φ^k^* randomly chosen from the underlying network.

We applied least squares polynomial curve fitting^36^ to fit the LCC z-scores best to their fold-change cutoffs, formulated by a function:

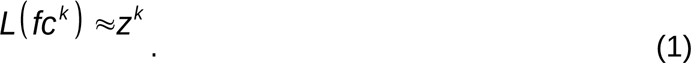

Our framework uses this function that helps in discovering the two peaks (*L*) patterns, the first at *fc^low^* and the second at *fc^high^*. The two local maxima, *L*(*fc^low^*) and *L*(*fc^high^*), identify two LCCs induced by the sets *Φ^low^* and *Φ^high^*, respectively.

The LCC given by the first peak, at *L*(*fc^low^*), represents the disease neighborhood. The LCC induced by *Φ^high^* is smaller than *Φ^low^*, but the DEGs at this threshold have a higher fold-change cutoff (*fc^high^)*. (Details in SM Note 1.4).

### Identification of peripheral and core regions

To identify the peripheral and core regions of each disease, the main criterion was to detect the two distinct local maximum peaks with least squares polynomial fitting curve (see SM Note 1.5 for details). To achieve this, we defined the *d_ratio_*, which quantifies the amplitude ratio of the second local maximum peak at *fc^high^* of *L*: *d_ratio_=z^amplitude^/z^max^*, where *z^max^* is the maximum value of *L*, *z^amplitude^= z^high^-z^vally^* with *z^high^* being the LCC z-score of *L*(*fc^high^*) and *z^vally^* denotes the minimum z-score between the two local maximum peaks *L*(*fc^low^*) and *L*(*fc^high^*) as illustrated in Fig. S1a.

Further, we applied criteria to decide the core and peripheral regions: (1) the LCC z-score induced by the core genes need to be larger than 1.6, (2) the LCC z-score induced by the peripheral genes need to be greater than 1.6, and (3) *d_ratio_*>0.1 (the average *d_ratio_* for 100 randomized data sets was 0.09). From the total 184 data sets, we found that 135 (73.4%) data sets fulfilled the above criteria for 104 distinct diseases. We selected 1.6 as a z-score threshold based on previous suggestion for identifying the significant connected component^5^. Topological and biological details of the core and peripheral patterns for 104 diseases are detailed in Supplementary Table 5. Full details on data sources, processing, and analysis are provided in SM Note 1.

### Robustness analysis

To validate the robustness of the pattern, we used two perturbation strategies: (1) detecting the pattern in different types of protein-protein interaction (PPI) networks, (2) changing the labels of disease and healthy samples randomly. Full details on data sources, processing, and analysis are provided in SM Note 2.

### Topological characteristics analysis

To show the topological differences between peripheral and core genes, we calculated the degree, betweenness^17,46^, closeness^17^, and clustering coefficient^18^ centralities in the human interactome. We used the Wilcoxon signed-rank test^47^ to calculate the p-value. A core is topologically different from a periphery if the Wilcoxon signed-rank test between the two distributions of centrality values (core genes and peripheral genes) is lower than or equal to 0.05 (threshold for which the two distributions are statistically significantly different).

To study the robustness of the disease neighborhood we considered its stability by fragmentation analysis: We defined immediate neighbors of the core in a disease neighborhood as the Dominate Range (DR) and measured fragmentation of the disease neighborhood by the size of the largest remaining connected component (LRCC) after removing the core and its DR. To measure the significance of the stability of the neighborhood we compared the size of the LRCC to 1,000 random counterparts. Furthermore, we analyzed the cohesiveness of the core to study to which extent the core corresponds to a topological cluster. A cluster represents a locally dense sub-graph that has more internal than external connections^48^, while on the contrary, a stretched sub-graph has significantly less internal connection. Therefore, we define the cohesiveness value (CV) of a given core *CO* as:

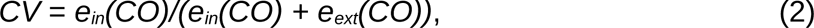

where *e_in_*(*CO*) denotes the number of internal edges and *e_ext_*(*CO*) denotes the external edges of the core (Fig. 3d). We generated 1,000 random connected components (RCCs) from the periphery as random counterparts to uncover the significant low cohesiveness of the core. To examine the significance of these observed values we computed their corresponding z-scores: (1) DR z-score,(2) core-removed z-score, (3) DR-removed z-score, and (4) Cohesiveness z-score (see SM Note 3 for full details).

### Hypergeometric test

The hypergeometric distribution gives the probability of k successes in n draws (without replacement) from a total population of size N, where K objects are considered as being a success. The probability that a drawn sample consists of k successes can be computed using:

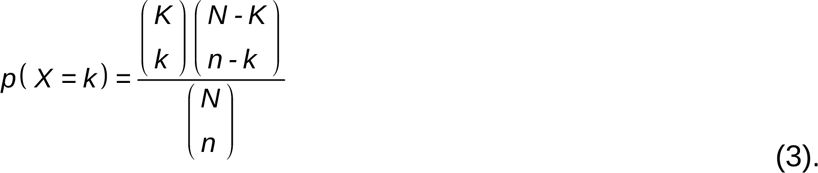

where X is a random variable. The corresponding p-value can then be computed using the survival function given by the following formula:

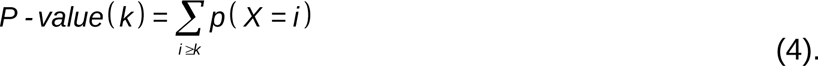

To evaluate whether or not the overlap between two different sets of core genes X and Y is higher than expected, we set the parameters of equation (3) to *n* = |X|, *k*= |*X*∩*Y*|, *K* = |Y|, and *N* equals to the total number of genes in the human interactome.

### Biological significance analysis

To evaulate the biological differences between peripheral and core genes, we collected 280 disease-causing intrinsically disordered proteins (IDPs) from the DisProt Database^21^ and 6,464 disease genes annotated by GWAS and OMIM. Next, we constructed a unique global core and a unique global periphery by combining the results from the 135 data sets. Next, we used the hypergeometric distribution to compute a p-value for measuring the statistical significance. We tested the enrichment of IDPs and disease genes for (1) global core, and (2) global periphery. As an example, we showed the biological relationship between the four sets for asthma GSE65204 (see SM Fig. S5a and SM Note 4.2 for full details).

### The key role of the core in disease pathways

To explore the role of specific cores in key disease pathways, we gathered 20 cancer gene expression data and 11 different autoimmune diseases. For cancer, we selected the cancer pathway from KEGG database (KEGG^49^: hsa05200), which includes 397 cancer genes. For autoimmune disease, we selected vitamin D pathway form BioGRID^50^ and WikiPathways which is comprised of 211 related genes (disease and gene lists can be found in Supplementary Table 6). Further, we used the hypergeometric p-value to determine the significance of the enrichment of the cores and the first neighbours of the core (DRs) in the cancer pathway and vitamin D pathway respectively (see SM Note 5 and Fig. 4c for full details).

### Quantification of disease-disease relationships

#### Construction of the disease-disease network

We used the Jaccard index, Sørensen–Dice coefficient, Simpson index, cosine index, and geometric index^35^ to quantify the proportion of shared genes between two gene sets. As an example, for each disease pair (*i, k*), based on their peripheral regions *PE_i_* and *PE_k_*, the overlap was measured by the Jaccard index *J*= |*PE_i_*∩*PE_k_* |/|*PE_i_*∪*PE_k_* |. The value lies in the range of [0, 1], where *J* = 0 if *PE_i_* and *PE_k_* have no genes in common and *J* = 1 indicates identical peripheral regions. Further, we calculated the *P-value_i,k_* based on the hypergeometric distribution to evaluate whether the overlap between two disease peripheries (Jaccard index *J)* are higher than expected in the human interactome (SM Note 6).

Furthermore, to quantify the disease-disease relationships, we defined the overlap peripherial similarity between two diseases *i* and *k* as:

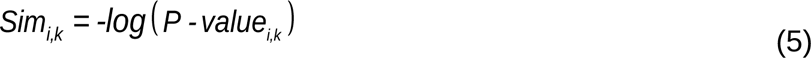

Next, we quantified the connection specificity index (CSI)^35^ of diseases based on the peripherial. The CSI is the number of diseases whose periphery is overlapping with the peripheries of i and k (Sim*_i,j_* and Sim*_k,j_*) which have a smaller overlap than the peripheries of i and k (Sim*_i,j_ < Sim_i,k_ and Sim_k,j_ < Sim_i,k_*), divided by the number of all diseases:

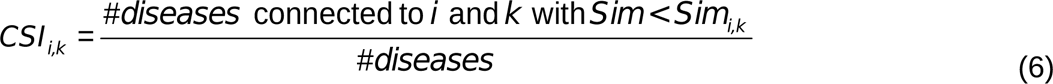

We used a cutoff of CSI>0.1 to define the lower boundary of interaction links. All the 104 diseases were mapped to 16 MeSH categories manually (details can be found in Supplementary Table 7, see Fig. 6a for an illustration).

#### Application in drug repurposing

For drug repurposing among similar diseases based on overlapping peripheries, we used 6,677 approved drug-disease pairs from the repoDB database (http://apps.chiragjpgroup.org/repoDB/), which includes 1,571 drugs (mapped to drug labels^51,52^) and 2,051 United Medical Language System (UMLS) disease concepts. From repoDB, we curated 384 drugs for 61 diseases that resulted in 704 approved drug-disease pairs. Next, we collected the gene targets of these 384 drugs in DrugBank database (Version 5.0, https://www.drugbank.ca/), the results can be found in the Supplementary Table 8. Finally, for each disease pair, we evaluated whether the overlapping part of their peripheries has more common drug targets than expected. We used a hypergeometric distribution to calculate the p-value for enrichment, with the null hypothesis that common drug targets are randomly drawn from all genes in the human interactome (see SM Note 6 for full details).

## Acknowledgments

This work was supported by the NSFC (No. 61772395, 61532014, 61672406 &61672407) and Shanghai Municipal Science and Technology Major Project (No.2018SHZDZX01), LCNBI and ZJLab. We acknowledge support from the U.S. National Institutes of Health (NIH) through the following grants: R01 HL118455, P01 HL13285, and K01 HL127265. The funders had no role in study design, data collection and analysis, decision to publish, or preparation of the manuscript.

## Author contributions

B.W. and A.S. conceived the project and wrote the manuscript. B.W. implemented the software and performed the main analyses. M.S., A.R., K.G., D.C.C.-C. and B.A.R. helped in writing the manuscript. A.S. supervised the project. S.T.W. provided supervision and funding for the project.

## Additional information

Competing financial interests: The authors declare no competing financial interests.

